# Transient cell-in-cell formation underlies tumor resistance to immunotherapy

**DOI:** 10.1101/2020.09.10.287441

**Authors:** Amit Gutwillig, Nadine Santana-Magal, Leen Farhat-Younis, Diana Rasoulouniriana, Asaf Madi, Chen Luxenburg, Jonathan Cohen, Krishnanand Padmanabhan, Noam Shomron, Guy Shapira, Meora Feinmesser, Oran Zlotnik, Reno Debets, Nathan E. Reticker-Flynn, Peleg Rider, Yaron Carmi

## Abstract

Despite the remarkable success of immunotherapy in cancer, most patients will develop resistant tumors. While the main conceptual paradigm suggests that relapsed clones emerge through a process of clonal selection and immunoediting, currently little evidence directly demonstrates this process in epithelial cancers.

To study this process, we established several mouse models in which tumors drastically regress following immunotherapy, yet resistant tumors relapse within a few weeks of treatment cessation. Whole exome analyses indicated that relapsed tumors share hundreds of neo-antigens with the primary tumors and are comparably killed by reactive T cells. Examination of tumor cells that survive immunotherapies revealed that they structure a transient cell-in-cell formation, which is impenetrable to immune-derived cytotoxic compounds and to chemotherapies. This formation is mediated predominantly by a cell-membrane protein on activated T cells, which subsequently induces epidermal growth factor receptors and STAT3 phosphorylation in tumors cells. In contrast to previous reports on cell-in-cell formations, here both cells remain alive and can disseminate into single tumor cells once T cells are no longer present.

Overall, this work highlights a powerful resistance mechanism which enable tumor cells to survive immune pressure and provides a new theoretical framework for combining chemotherapies and immunotherapies.

## Main

Given the direct correlation between the prevalence of cytotoxic T-cell immunity and improved clinical outcomes, and survival^1–3^, most therapeutic strategies are aimed at harnessing T-cell immunity to fight cancer. These attempts include *de novo* expansion of tumor-infiltrating cytotoxic T^4,5^, engineered T cells^6,7^, or blocking antibodies directed against suppressive receptors^8,9^. However, long-term follow-up indicates that while patients will experience tumor regression, it is frequently followed by recurrence of tumors that are largely resistant to subsequent treatments^10–12^. As tumor cells often express altered or neo-antigens^13,14^, it remains unclear why eliciting the full spectrum of T-cell immunity to these antigens is not sufficient to eradicate tumors.

Two theoretical frameworks have been widely employed to understand how tumors escape T-cell-based therapies. The main conceptual paradigm suggests that in order to avoid killing by T cells, tumor cells edit or lose their targeted antigens, or down-regulate HLA molecules^15–17^. Alternately, tumor resistance to immunotherapy was shown to be acquired through tumor cell-intrinsic mechanisms, including loss-of-function of JNK and PTEN, MYC overexpression, and constitutive WNT signalling^18–20^.

Nonetheless, most of our knowledge on this process comes from experiments comparing individuals who respond or do not respond to immunotherapy and not from experimental settings that measures the cell-intrinsic changes, neo-antigen landscape, and immune parameters within the same individual. Furthermore, whole exosome and genome sequencing comparisons of paired primary and relapsed tumors indicate that the majority of clones in relapsed tumors are shared with the primary tumors, suggesting relative low rates of clonal deletion^21–23^.

Therefore, to study this process in melanoma and breast cancer, we developed several mouse models in which resistant tumors relapse following immunotherapies. In both mouse and humans, we found that the tumor cells remaining after immunotherapy form unique cell-in-cell structures and generate membrane architecture that is impenetrable by immune-derived lytic granules, cytotoxic compounds, and chemotherapies. Both cells in this formation maintain their integrity and viability and can survive for weeks in culture containing reactive T cells. Once T cells are removed, tumor cells disseminate back into their parental single cells. This biological process has never been described before and is the main mechanism through which tumor cells escape T-cell immunity and give rise to relapsed tumors.

## Results

### Relapsed tumor cells share neoantigens with primary tumors and are equally susceptible to killing by T cell

We have recently reported that the combination of dendritic cell adjuvant with tumor-binding antibodies elicits an extremely potent T-cell reaction^24^. Indeed, when palpable melanoma tumors were injected with a combination of TNFα, CD40L, and an antibody against the melanoma antigen TRP1, tumors were almost completely eradicated in all mice. Nonetheless, after about ten days, about half the mice developed reoccurred tumors that were resistant to subsequent treatments (Fig. 1a). Similar patterns of tumor relapse and resistance were also observed upon treating BALB/c mice bearing breast tumors treated with allogeneic antibodies and DC adjuvant (Fig. 1a). Intriguingly, immune composition analysis of tumors following the second immunotherapy indicated treated mice across all immunotherapies show massive immune-cell and T-cell infiltration (Fig. 1b, Extended Data Fig. 1a) with a dominant representation of T-cell clones against TRP2 and gp100 (Fig. 1c). Furthermore, analysis of melanoma cells showed they maintain expression of MHC-I and MHC-II (Fig. 1d, Extended Data Fig. 1a–1b), as well as TRP2 and gp100 (Fig. 1e). To assess the overall changes in immunogenicity of resistant cell lines, beyond TRP2 and gp100 expression, we analysed their neoantigen landscape in comparison to that of B16F10 from untreated mice (Fig. 1f). Whole exome analysis (WES) indicated comparable neoantigen burden across all samples (Fig. 1g) with the same 25 germline mutations in the highest score ones (Fig. 1h, Supplementary Table-1). RNAseq analyses further demonstrated similar expression levels of known B16F10 neoantigens in B16F10 cell lines established from relapsed tumors as well as in tumors cells sorted from mice following effective immunotherapy (Fig. 1i, Supplementary Table-2).

**Fig. 1:**
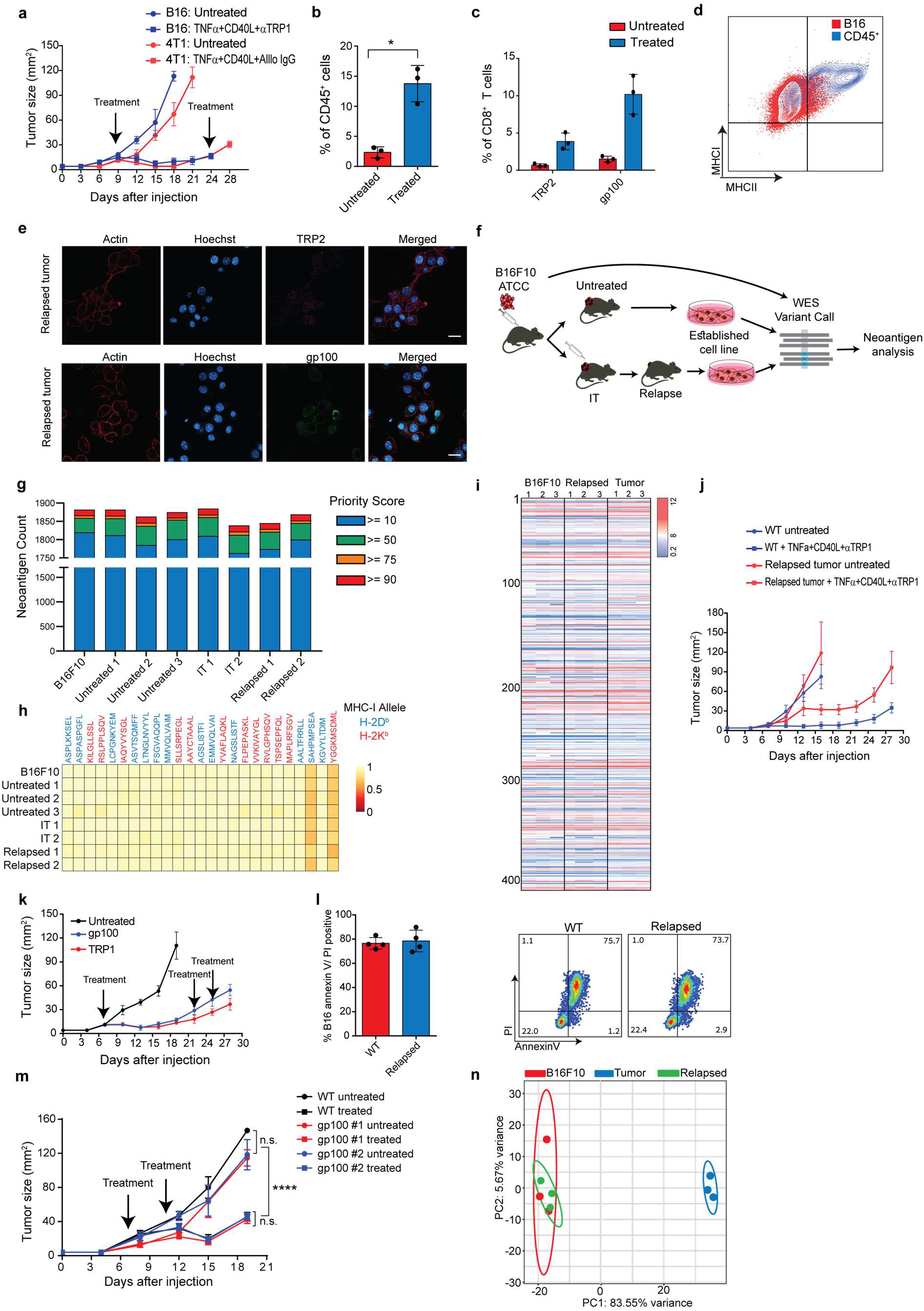
Relapsed tumor cells share neoantigens with primary tumors and are equally susceptible to killing by T cells. **a.** Tumor size following treatments with anti-CD40, TNFα, and tumor-binding antibodies (n=4). **b.** Mean percentages of T cells out of CD45^+^ cells in relapsed tumors (n=4). **c.** Mean percentage of TRP2- and gp100-reactive T-cell clones in relapsed tumors (n=4). **d.** Representative Flow cytometry of MHC-I and MHC-II expression in relapsed B16F10 cells. **e.** Representative staining of TRP2 and gp100 in relapsed B16F10 cells. **f.** Illustration of neoantigen discovery pipeline. **g.** Neoantigen burden in B16F10 cells isolated from untreated tumors, or following immunotherapy treated tumors (IT). **h.** Allele frequency comparison of 25 neoantigens with the highest MHC-I affinity. **i.** RNAseq expression level of B16F10-known neoantigens. **j.** Tumor growth following treatments with anti-CD40, TNFα, and anti-TRP1 tumor-binding antibodies (n=4). **k.** Tumor size following adoptive transfer of gp100-reactive CD8^+^ or TRP1-reactive CD4^+^ T cell (n=4). **l.** Mean percentages of apoptotic tumor cells after incubation overnight with gp100-reactive CD8^+^ T cells (n=5). **m.** Tumor size following adoptive transfer of gp100-reactive T cell (n=4). **n.** Principal Component Analysis (PCA) of B16F10 tumor cell lines freshly isolated from treated mice. Experiments were repeated independently at least three times. Statistical significance was calculated using ANOVA with Tukey’s correction for multiple comparisons (*** denotes p<0.001, **** denotes p<0.0001). Error bars represent standard error. Scale bars = 20 μm.

To assess whether tumors acquire resistance through inherent mechanisms, we next tested the susceptibility of these cell lines to immunotherapy. Consistent with their patterns of gene and neoantigens expression, these cell lines were equally susceptible to immunotherapy as their parental B16F10 cells (Fig. 1j). To test if this occurrence represents a broader phenomenon, we also treated melanoma-bearing mice with splenic CD8^+^ T cells expressing TCR against gp100, or TRP1 melanoma antigens. Both treatments induced significant tumor regression, followed by tumor reoccurrence in all treated mice (Fig. 1k). As observed with the previous immunotherapy, reoccurred tumors were resistant to subsequent treatment with gp100- or TRP2-reactive T cells (Fig. 1k). Consistently, *in vitro* analysis indicated that cell lines established from resistant tumors were susceptible to killing by gp100-reactive T cells (Fig. 1l). Moreover, mice bearing these cell lines were equally susceptible to treatment with gp100-reactive T cells as mice bearing the parental melanoma cells (Fig. 1m).

These results suggest that transient *in vivo* mechanisms rather than inherited ones govern resistance of relapsed tumors. In support of this hypothesis, RNAseq analysis indicated that of all established cell lines cluster within the same principle components, whereas the expression profile of freshly sorted tumor cells from immunotherapy-treated mice is markedly distinct (Fig. 1n, Extended Data Fig 1c, Supplementary Table-3).

### Tumor cells remaining after T-cell killing organize in a cell-in-cell formation

To better characterize the tumor cells that survive in mice following immunotherapy, we enzymatically digested treated tumors and sorted the live melanoma cells. Confocal analyses indicated that most of the tumor cells organize in clusters of several nuclei surrounded by a single membrane and cortical actin, suggesting they structure in a cell-in-cell formation (Fig. 2a). Transmitting electron microscopy (TEM) of freshly sorted cells further corroborated a distinct segregation of the membranes and cytosols of the two cells. The inner cell was dense and seems to have been compact within another cell (Fig. 2b). To ensure that these structures do not result from the isolation procedure, we also analysed histological sections of tumors, whose nucleus and cell membranes are fluorescently labelled. While untreated tumors showed low levels of cell-in-cell clusters, about half the cells that survived in treated mice were organized as such (Fig. 2c–2d).

**Fig. 2:**
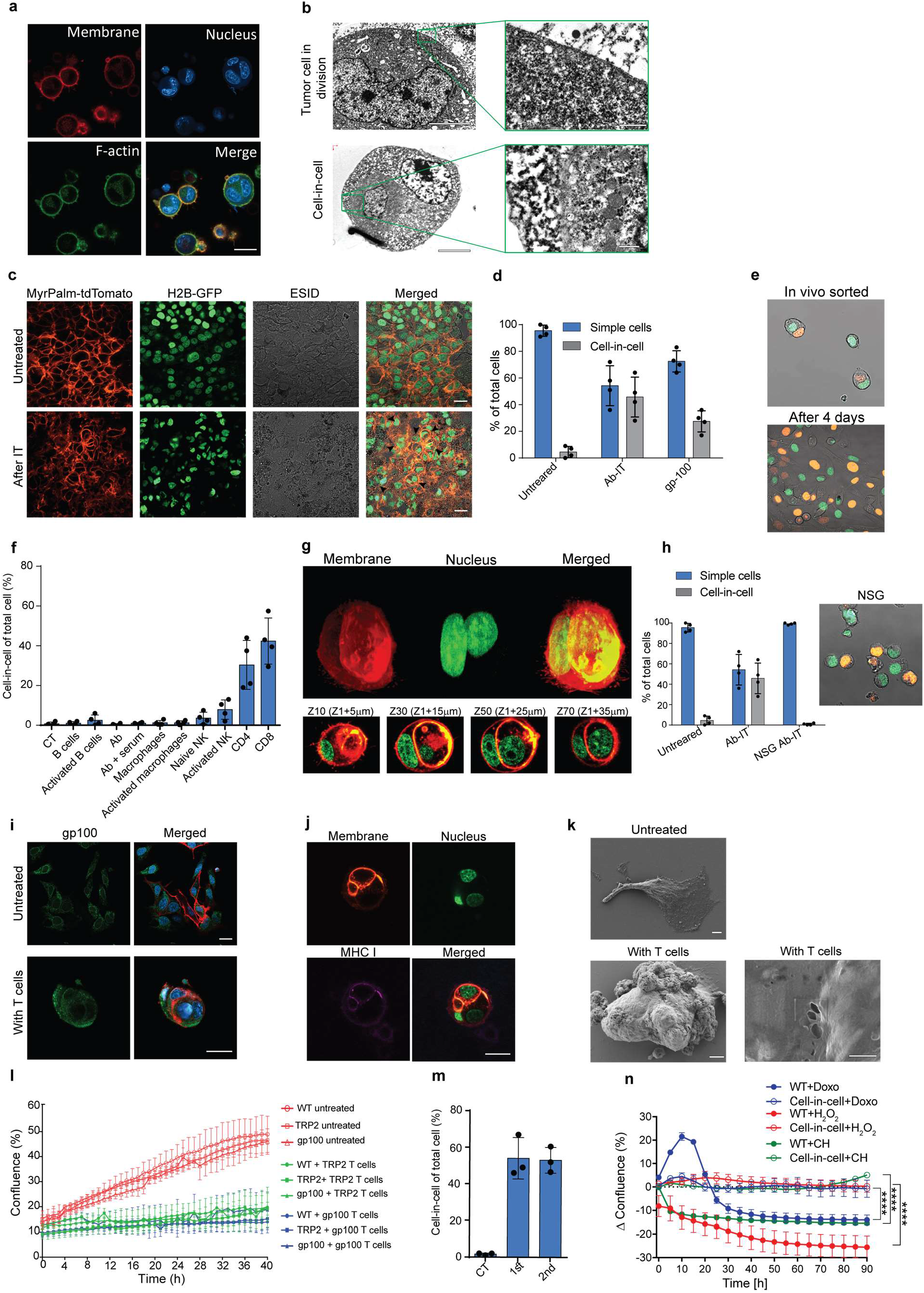
Tumor cells remaining after T-cell killing organize in a cell-in-cell formation. **a.** Representative images of B16F10 tumor cells sorted from tumors relapsed after immunotherapy. **b.** Transmitting electron microscopy images of relapsed B16F10 tumor cells. **c.** Histological sections of B16F10 tumor untreated, and five days following immunotherapy. **d.** Mean percentage of cell-in-cell and single cells in untreated B16F10 tumor-bearing mice, treated with anti-CD40, TNFα, and anti-TRP1 (Ab-IT), or treated with gp100-reactive (gp100 ACT) (n=4). **e.** Representative images of B16F10 labeled with H2B-tdTomato and H2B-GFP freshly isolated from tumor-bearing mice treated anti-CD40, TNFα, and anti-TRP1. **f.** Mean percentage of cell-in-cell formation following incubation of B16F10 cells with immune cells (n=3). **g.** Representative 3D projection and horizontal sections (Z-stack) of B16F10 cells incubated with gp100-reactive CD8^+^ T cells. **h.** Mean percentage of cell-in-cell and single cells in tumor-bearing NSG^-/-^ mice treated with anti-CD40, TNFα, and anti-TRP1 antibodies (n=3). **i.** Representative staining of gp100 expression in B16F10 tumor cells following incubation with gp100-reactive CD8^+^ T cells. **j.** Representative images of MHC-I expression in B16F10 tumor cells following incubation with gp100-reactive CD8^+^ T cells. **k.** SEM analysis of B16F10 incubated with gp100-reactive CD8^+^ T cells. **l.** Percentages of B16F10confluence tumor cells following incubation with gp100- and Trp2-reactive T cells (n=3). **m.** Mean percentage of cellin-cell in B16F10 culture following one (1^st^) or two (2^nd^) cycles of incubation with gp100-reactive CD8^+^ T cells (n=3). **n.** Mean percentage of confluence change over time of B16F10 control (WT) or cell-in-cell tumors (n=3). Experiments were repeated independently at least three times. Statistical significance was calculated using ANOVA with Tukey’s correction for multiple comparisons (*** denotes p<0.001, **** denotes p<0.0001). Error bars represent standard error. Error bars represent standard error. Scale bars = 20 μm (**a, c, I, j**), 5 μm (**b** left, **k** left), 500 nm (**b** right, **k** right).

To test if this formation was merely the result of incomplete cell division, we injected into mice tumor cells whose nuclei are labelled with GFP or with tdTomato, and treated the mice with immunotherapy. About one third of the tumor cells were a mixture of both GFP and tdTomato nuclei (Fig. 2e). Furthermore, monitoring the dynamics of single cells sorted from treated mice indicated that they spontaneously clustered in a cell-in-cell formation during their *in vitro* culture (Supplementary Movie-1). Over time, all cells disseminated into single cells bearing morphological features similar to those of the parental cell line with no observed cells concomitantly expressing the two fluorophores (Fig. 2e, Supplementary Movie-2). To test whether a specific immune cell type mediate these cell-in-cell formations, we isolated the main effector cells from tumor-bearing mice and incubated them overnight with B16F10 tumor cells. We found that only T cells—mainly CD8^+^, and with slower kinetics also CD4^+^—induce a cellin-cell formation with the same special characteristics as the ones observed *in vivo* following immunotherapy (Fig. 2f–2g, Extended Data Fig. 2a–2b, Supplementary Movies-3). Consistently, no cell-in-cell structures were observed in Nude SCID-gamma^-/-^ (NSG) mice treated with immunotherapy (Fig. 2h).

We next tested whether this phenomenon occurs in other tumor cell types. Indeed, the same structures were also observed in 4T1 mammary carcinoma incubated with allogeneic T cell, but not in immortalized mammary epithelial cells (Extended Data Fig. 2c–2d). Most importantly, similar structures were also found in human breast carcinoma and melanoma cell lines incubated with allogeneic T cells from healthy donors (Extended Data Fig. 2e–2f). Furthermore, incubation of cell lines established from two melanoma patients with their corresponding TIL indicated that the vast majority of cells that survive in culture form a cellin-cell formation, similar to the one found in mice (Extended Data Fig. 2g).

We next assessed how this structure protects the tumor cells. Initially, we incubated tumor cells with T cells expressing TCR against the known melanoma antigen gp100 and tested whether they escape T cells by reducing their MHCI or gp100 expression. We found that all cells in the cluster expressed similar levels of MHCI or gp100 compared to untreated tumor cells (Fig 2i–2j). Furthermore, scanning electron microscope (SEM) analysis indicated that multiple T cell were attached to the clustered tumor cells, and many pores were observed on the outer cell membrane, suggesting their ability to form an immunological synapse on their membrane (Fig. 2k). Next, we tested if tumor cell lines established following killing with TRP2- and gp100-reactive T cells would be less sensitive to killing by the same reactive T cells. Thus, cell lines were then incubated again with gp100 and TRP2-reactive T cells, and their rates of killing were compared to that of the parental cell line (Fig 2l). Instead, tumor cells re-organized in cell-in-cell formation regardless of the nature of the tumor-reactive T cells (Fig. 2m), suggesting that this formation provides a transient protection, rather than an inherited one. Thus, we compared the sensitivity of single-tumor cells and those in a cell-in-cell formation to a variety of cytotoxic compounds and chemotherapies. In all the tested compounds, cell-in-cell formations survived longer and higher concentrations compared to their parental cell lines (Fig. 2n and Extended Data Fig. 2h). To visualize the penetration rates of cytotoxic compound to these structures, we incubated CFSE-labelled lysosomes with single-tumor cells with cell-incell formed tumor cells. Indeed, lysosome penetration accumulated on the multiple membranes of the cell-in-cell formations, and penetration of the latter was much slower compared to single-tumor cells (Extended Data Fig. 2i).

### IFNγ induces membrane-bound protein on T cells that mediate tumor cell-in-cell formations

To understand why only T cells induce a cell-in-cell formation, we initially analysed their interactions with tumor cells using SEM. Interestingly, tumor-reactive T cells, but not PMA plus ionomycin-activated NK cells or LPS-activated macrophages, express the intact form of lysosomes on their cell surface (Fig. 3a–3c, Extended Data Fig. 3a). Next, we tested if this formation can be induced by other extracellular vesicles. Hence, B16F10 cells were cultured overnight with extracellular vesicles from activated NK, macrophages, immortalized melanocytes (Melan A), B16F10-derived lysosomes, or commercial 1μM latex beads. Confocal analysis of these cultures indicated that only T cell-derived lysosomes induced a cell-in-cell formation, suggesting this it is mediated by T cell-restricted molecules (Fig 3d–3f).

**Fig. 3:**
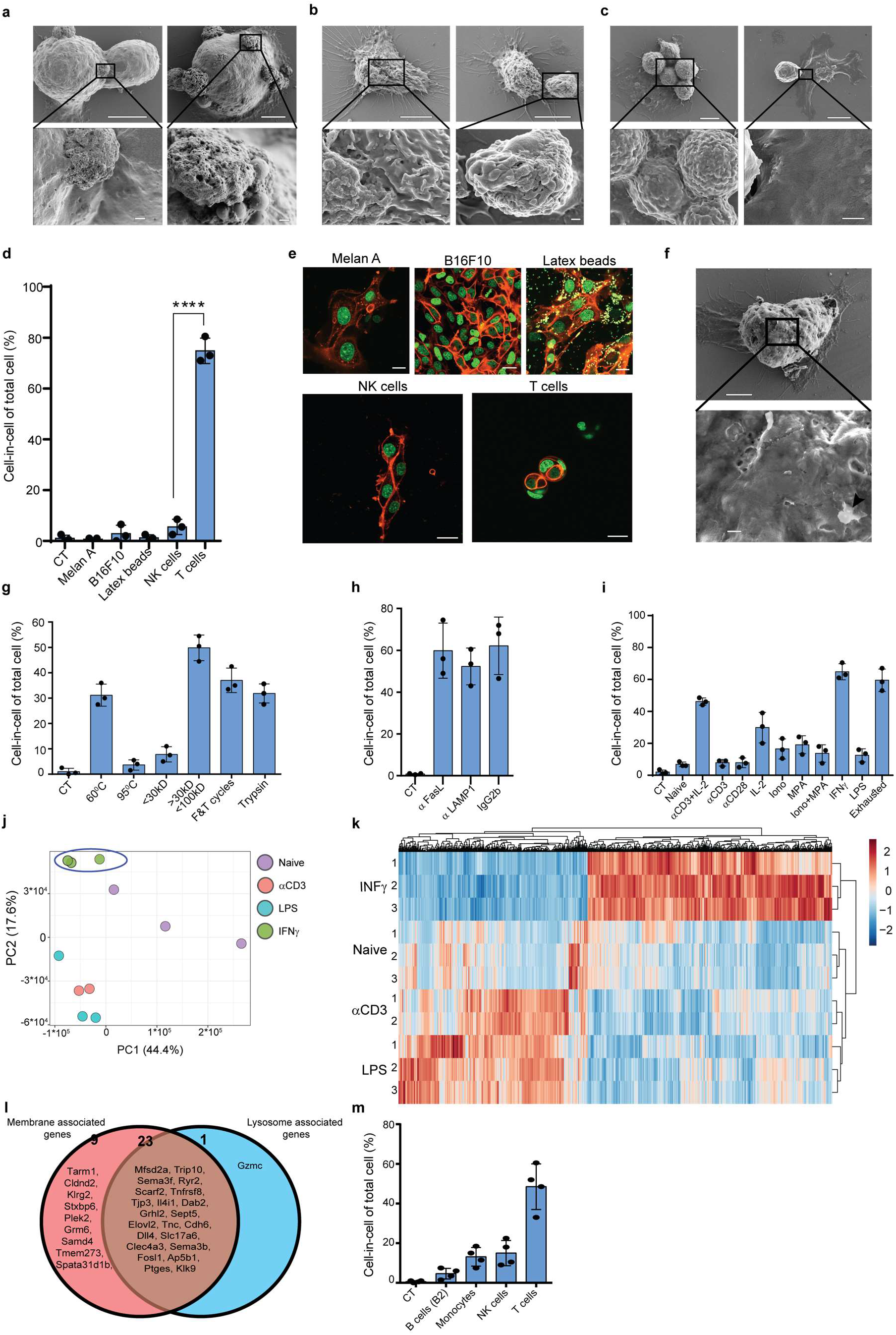
IFNγ induces membrane-bound proteins on T cells that mediate tumor cell-in-cell formation. **a.** SEM analysis of B16F10 incubated with gp100-reactive CD8^+^ T cells. **b.** SEM analysis of B16F10 incubated with peritoneal macrophages. **c.** SEM analysis of B16F10 incubated with NK cells. **d.** Mean percentage of cell-in-cell of B16F10 incubated overnight with latex beads, B16F10 and Melan A melanosomes, NK- and T cell-derived lysosomes (n=3). **e.** Representative images of B16F10 incubated overnight with latex beads, B16F10 and Melan A melanosomes, NK- and T cell-derived lysosomes. **f.** SEM analysis of B16F10 incubated with T cells-derived lysosomes (arrowhead). **g.** Mean percentage of cell-in-cell formation after incubation with modified T-cell lysosomes (n=3). **h.** Mean percentage of cell-in-cell formation after incubation with T cells and lysosomes-blocking antibodies (n=3). **i.** Mean percentage of cell-in-cell formation following incubation with pre-activated T cells (n=3). **j.** PCA of expression profiles of naïve and activated CD8^+^ T cells **k.** Heatmap of gene expression patterns of naïve and activated CD8^+^ T cells. **l.** Venn diagram of membrane- and lysosome-associated genes up-regulated in CD8^+^ T cells stimulated with IFNγ. **m.** Mean percentage of cell-in-cell formation following overnight incubation with IFNγ-activated immune cells. Experiments were repeated independently at least three times. Statistical significance was calculated using ANOVA with Tukey’s correction for multiple comparisons (*** denotes p<0.001, **** denotes p<0.0001). Error bars represent standard. Scale bars = 5 μm (**a-c** top, **f** top), 500 nm (**a-c** bottom, **f** bottom), 20 μm (**e**).

To shed light on the nature of these molecules, we isolated the lysosomes from the supernatants of activated T cells and subjected them to biochemical manipulations before adding them to a B16F10 culture. Boiling the lysosomes at 95°C for 5 min completely abrogated their capacity to induce a cell-in-cell formation, but neither incubation at 60°C for an hour nor freeze-thaw cycles abrogated this capacity, suggesting that these are globular proteins and not enzymes (Fig. 3g, Extended Data Fig. 3b). Size exclusion assays indicated that these proteins are larger than 30kDa, smaller than 100kDa, and are not digested by trypsin (Fig. 3g, Extended Data Fig. 3b). Nonetheless, incubation of tumor cells with gp100-reactive T cells in the presence of blocking antibodies against lysosome-associate proteins such as LAMP1, CD95, or CD63 did not alter their capacity to induce a cell-in-cell formation, suggesting that these molecules are also expressed on T-cell membranes (Fig. 3h). To elucidate the type of activations that induce these molecules on T cells, we incubated naïve splenic T cells overnight with various stimulators, washed them, and incubated them with tumor cells. We found that T cells activated with IFNγ, and to a lesser extent also anti-CD3 plus high-dose IL-2, induce the most cell-in-cell formation (Fig. 3i, Extended Data Fig. 3c). We then compared the expression profile of IFNγ-activated T cells to that induced by anti-CD3 or LPS, which do not induce cell-in-cell formation (Fig 3j). Indeed, IFNγ-activated T cells exhibit a distinct expression pattern (Fig. 3k, Supplementary Table-4). Of these genes, only about 30 proteins are expressed on the cell membrane, and most of them are shared with lysosome membranes (Fig. 3l). We next tested if other IFNγ-activated cells can induce a cell-in-cell formation, or whether it is restricted to activated T cells. We found that IFNγ-stimulated NK cells, but not macrophages or B cells, could induce a tumor cell-in-cell formation, albeit at a lower percentage compared to activated T cells (Fig. 3m).

### Cell-in-cell tumor formation is governed by STAT3 signalling

We then set out to understand how tumor cells form cell-in-cell structures and what mechanisms govern this structure. Initially, we tested whether it is cell fusion. For that, we incubated tumor cells whose cytosols are labelled with either GFP or mCherry with tumor-reactive T cells. Confocal analysis indicated that each cell type in the cell-in-cell formation maintained its cytoplasm, and no mix between colours was detected (Fig. 4a). Furthermore, long-term followup of cell dissemination from this structure indicated that each cell maintains its initial single labelling colour (Extended Data Movie-4). To corroborate this, we also incubated tumor cells whose membrane, nucleus, and F-actin were labelled with different fluorophores. Similarly, each cell in this formation was separated and maintained the integrity of its original cell components (Fig 4b). We next tested the possibility that this formation is entosis by staining for molecules reported to mediate that structure. However, we observed no increase or changes in the cellular localization of phosphorylated β catenin, E-cadherin, and phosphorylated integrin βl, which are all important mediators of entosis^25^ (Fig. 4c–4e and Extended data Fig. 4a–4d). Furthermore, we observed no reduction in T cell-mediated cell-in-cell formation upon blocking of E- and N-cadherins, actin filaments disruption, or inhibition of Wnt signalling (Fig. 4f–4g). In sharp contrast, the blocking of mRNA synthesis or protein production completely abrogated tumor cells’ capacity to form a cell-in-cell formation, suggesting that the structure requires *de novo* synthesis of genes (Fig. 4f–4g). To assess what genes govern this formation, we compared the gene signature of untreated B16F10 cells, to cell-in cell formation induced following incubation with T cell-derived lysosomes and to that of tumor cells sorted from *in vivo* five days after immunotherapy. Over 600 genes were increased in cell-in-cell induced by T cell-derived lysosomes compared untreated B16F10 cells, all of which were also upregulated by the tumor cells sorted from treated animals (Fig. 4h, Supplementary Tables 5-6). Over 400 of these genes were part of the same pathway, highlighting a potential role for Jak/STAT3 and its upstream receptors such as EGFR and EGR1 (Fig. 4h). Consistent with mRNA levels, both cells with the cell-in-cell formation expressed higher levels of phosphorylated STAT3 (Fig. 4i, Extended Data Fig. 4e). Similarly, higher levels of phosphorylated EGFR1 were observed in cell-in cell tumors (Fig. 4j, Extended Data Fig. 4f). Importantly, inhibition of STAT3, or EGFR pathway, completely prevented cell-in-cell formation upon incubation with T cells or T cell-derived lysosomes (Fig. 4k–4l), corroborating the necessity of this pathway for the generation of cell-in-cell formation.

**Fig. 4:**
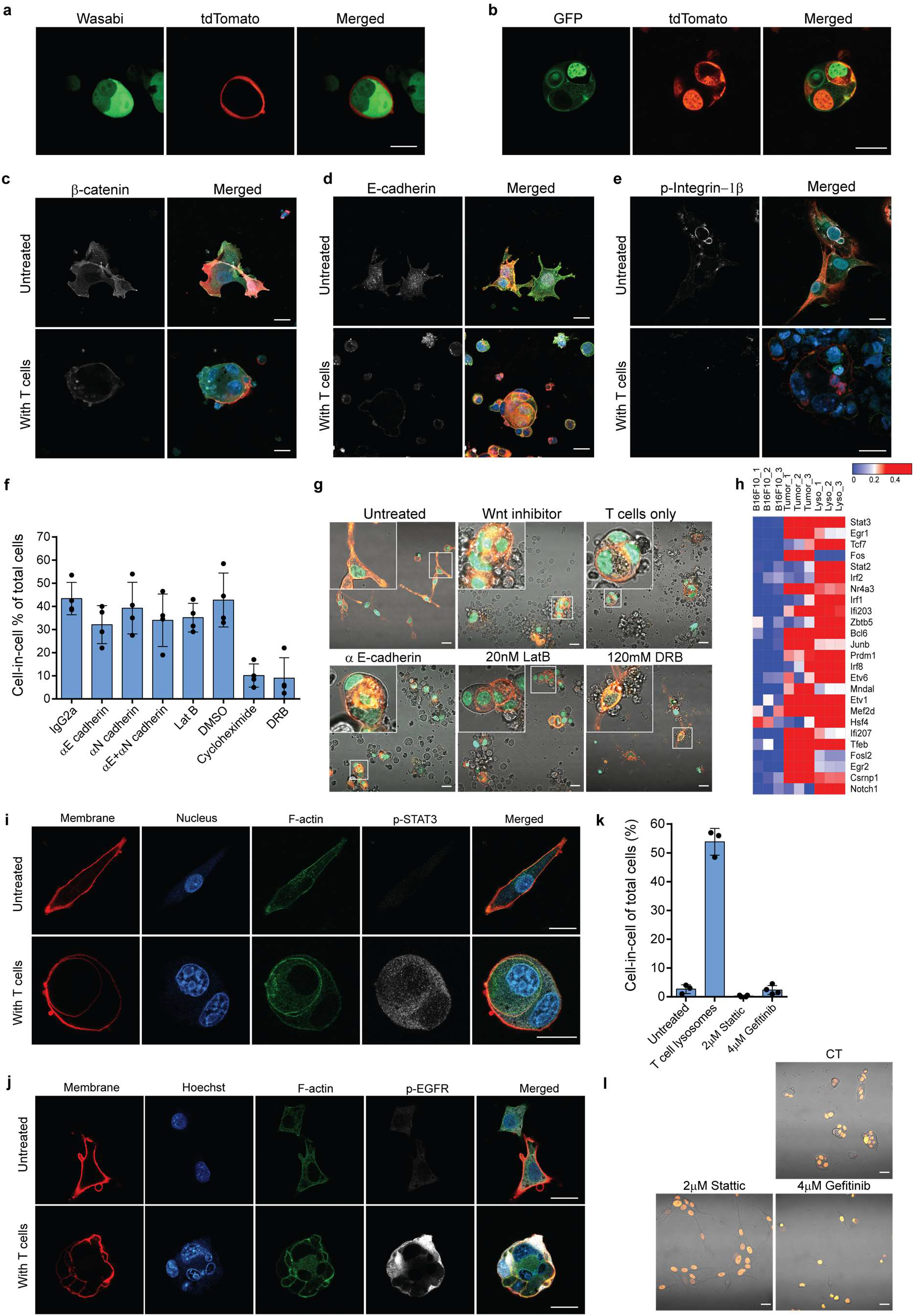
Cell-in-cell tumor formation is governed by STAT3 signalling and EGF receptors. **a.** Representative images of cytosol-labeled B16F10 cell lines incubated overnight with gp100-reactive T cells. **b.** Representative images of B16F10 cell lines labeled with H2B-GFP and MyrPalm-TdTomato, and B16F10 cell lines labeled with H2B-TdTomato and lifeactin-GFP, following overnight incubation with gp100-reactive T cells. **c.** Representative images of β-catenin expression in B16F10 cells incubated with gp100-reactive T cells. **d.** Representative images of E-cadherin expression in B16F10 incubated with gp100-reactive T cells. **e.** Representative images of phosphorylated integrin-1 β expression on B16F10 cells incubated with activated T cells. **f.** Mean percentage of cell-in-cell of B16F10 cells treated with different inhibitors and incubated with T cells (n=3). **g.** Representative images of B16F10 cells treated with inhibitors and incubated overnight with gp100-reactive T cells. **h.** Relative expression of highly enriched genes in B16F10 cells isolated directly from relapsed tumors (Tumor) and after incubation with T-cell lysosomes (Lyso), compared to B16F10 control cells (WT) (n=3). **i.** Representative images of phospho-STAT3 in B16F10 incubated with gp100-reactive T cells. **j.** Representative images of phosphorylated EGFR of B16F10 incubated with gp100-reactive T cells **k.** Mean percentages of cell-in-cell formation following overnight incubation with T cell-derived STAT3 or EGFR inhibitors (n=3). **l.** Representative confocal images of B16F10 cells incubated with STAT3 or EGFR inhibitors and with T cell-derived lysosomes. Experiments were repeated independently at least three times. Statistical significance was calculated using ANOVA with Tukey’s correction for multiple comparisons (*** denotes p<0.001, **** denotes p<0.0001). Error bars represent standard error. Scale bars = 20 μm.

## Discussion

Cancer immunoediting has been the primary paradigm explaining how tumors escape T-cell immunity and the framework for most recent immunotherapies^16,26^. In support of this view, fibrosarcoma clones emerged in T cell-deficient mice and were deleted upon injection to immunocompetent mice^27,28^. Immunoediting is further supported by clinical observations in which tumors relapse by omitting their expressed antigen^29–31^. While it is clear that immunogenic clones can be eliminated by T cells, the presence of immunogenic clones expressing neoantigens despite having infiltrated reactive T cells has been well documented^4,32,33^. Consistently, the persistence of TAA-reactive T cells correlates with excellent prognosis in responsive individuals^34,35^, indicating that immunogenic clones do not change so frequently. Furthermore, several seminal studies demonstrated that TCR-MHC is promiscuous, allowing each T cell to recognize several up to hundreds of different peptides^36–38^. As a result, editing a given antigen may not be sufficient to prevent T-cell recognition; or alternately, the antigen may be recognised by other T cells.

Whereas a suppressive microenvironment provides some explanation regarding how immunogenic clones can survive T-cell pressure^39–41^, why tumors reoccur in patients undergoing complete response remains unknown. Here, we found that tumor cells escape T-cell-based immunotherapies by forming a transient cell-in-cell structure, and not by immunoediting. These structures generate multiple membrane layers, thus reducing the probability of compounds’ internalization, hence leading to increased tumor-cell survival.

Several cell-in-cell formations have been described before including phagocytosis, cell cannibalism, and entosis^42^. Recently, these structures have been shown to enable tumor-cell survival after chemotherapy treatment by engulfing and digesting neighbouring cells^43^. Despite structural similarity to entosis, the cell-in-cell structure described in this study differs markedly in several key elements: First, it is triggered predominantly by reactive T cells, rather than glucose and nutrient starvation^44^, mitosis^45^, or loss of adherence^25^. Additionally, it requires *de novo* synthesis of genes regulated by STAT3, and not through Rho-Rock and β-cathenin signalling^25^. Most importantly, the structure described here is reversible and only rarely leads to cell death and apoptosis.

Overall, the ability of tumor cells to transiently enter and disseminate from each other in response to T-cell killing is a biological process that has never been described heretofore. It better explains how immunogenic tumors can survive in the host and provides a novel framework for immunotherapies.

## Material and Methods

### Tumor models

For melanoma tumor studies, 2.5×10^5^ B16F10 cells in 50 μL DMEM were injected s.c. into C57BL/6 mice above the right flank. For a triple-negative breast cancer model, 2× 10^5^ 4T1 cells in 30 μL DMEM were injected into mammary fat pad number five. Tumor size was measured twice a week using calipers. For immunotherapy, tumor-bearing mice were injected intratumorally twice, two days apart, with 100 μg anti-CD40 (FGK4.5; BioXCell), 0.2 μg TNFα (BioLegend), and 200 μg/mouse anti-TRP1 antibody (TA99; BioXCell). For adoptive cell transfer, splenic T cells were infected with pMIGII encoding TCR recognizing MHCI-gp100_25-33_^46^, or MHCI-TRP2_180-188_^47^, or MHCII-TRP1_113-126_^48^. Recipient mice were sub-lethally irradiated at a single dose of 600 rad, and injected i.v. twice, 3 days apart, with 1×10^6^ transduced T cells followed by i.p. injections of 300,000 IU of IL2 (PeproTech) for 4 consecutive days.

### Neoantigen predication

Genomic DNA was extracted from 5×10^6^ B16F10 tumor cell lines using NucleoSpin^®^ Tissue (MACHEREY-NAGEL), according to manufacturer instructions. Whole exome sequencing was performed by NovoGene. Sequencing libraries were constructed using the Agilent SureSelect kit and sequence on an Illumina Novaseq 6000 sequencer. Sequencing reads were filtered using the AGeNT SureCall Trimmer (Agilent, v4.0.1), and quality was evaluated using FastQC v0.11.3. Alignment to the mouse genome (mm10, GRCm38.87) was performed using the Burrows-Wheeler Aligner (bwa v 0.7.12) through the bwa mem command with the following parameters: -M -t 4 -T 30. SAMtools was used to remove alignments with a MAPQ score less than 5, and Picard was used to remove duplicate reads. Sorting and indexing were performed using SAMtools. Variants were called using MuTect2 (GATK v 4.0.0.0) and filtered using FilterMutectCalls. Variants were annotated with SnpEff. Neoantigens were identified using MuPeXI v1.2.0^49^ with the following parameters: -s mouse -a H-2-Db, H-2-Kb -l 9-11. MuPeXI was used in conjunction with NetMHCpan v.4.0 and Variant Effect Predictor (VEP) v.87.27. Heatmaps were generated in R using the “pheatmap” package (v1.0.12).

### Confocal microscopy

Cells were cultured on glass-bottom confocal plates (Cellvis). Cells were imaged live, or following fixation with 2% PFA for 20 minutes. Fixed cells were washed twice with PBS and stained for membranes with DiD membrane dye (Invitrogen), Alexa Fluor™ 594 Phalloidin (Invitrogen) and Hoechst 33342 (Fluka). For immunofluorescence, fixed cells were blocked with 5% BSA and mouse serum, stained overnight with primary antibodies, washed, and stained with secondary antibody. We used the primary antibodies: anti-CD44 (clone IM7, BioLegend), anti-MHC class I (clone 28-8-6, BioLegend), anti-β-catenin (clone D2U8Y, Cell Signaling Technology), anti-phosphorylated STAT3 (clone D3A7, Cell Signaling Technology), anti-active integrin 1β (Cell Signaling Technology), anti-integrin 1β (clone 9EG7), anti-E-cadherin (clone 36, BD Biosciences), anti-TRP2 (clone C-9, Santa Cruz Biotechnology), anti-gp100 (clone EP4863(2)), and anti-phosphorylated EGFR (clone EP38Y Abcam).

### Statistics

Significance was calculated using the nonparametric two-way ANOVA with Tukey’s correction for multiple hypotheses. In some cases, a Bonferroni-Sidak post-test was performed after two-way ANOVA. For two-group analysis, one-way ANOVA with Dunn’s test was performed. The results were analyzed by Prism (GraphPad Software, Inc.). All statistical analyses were performed in Prism (GraphPad Software, Inc.). All experiments were performed at least three times with at least three replications.

## Extended Figures

**Extended Data Fig. 1:**
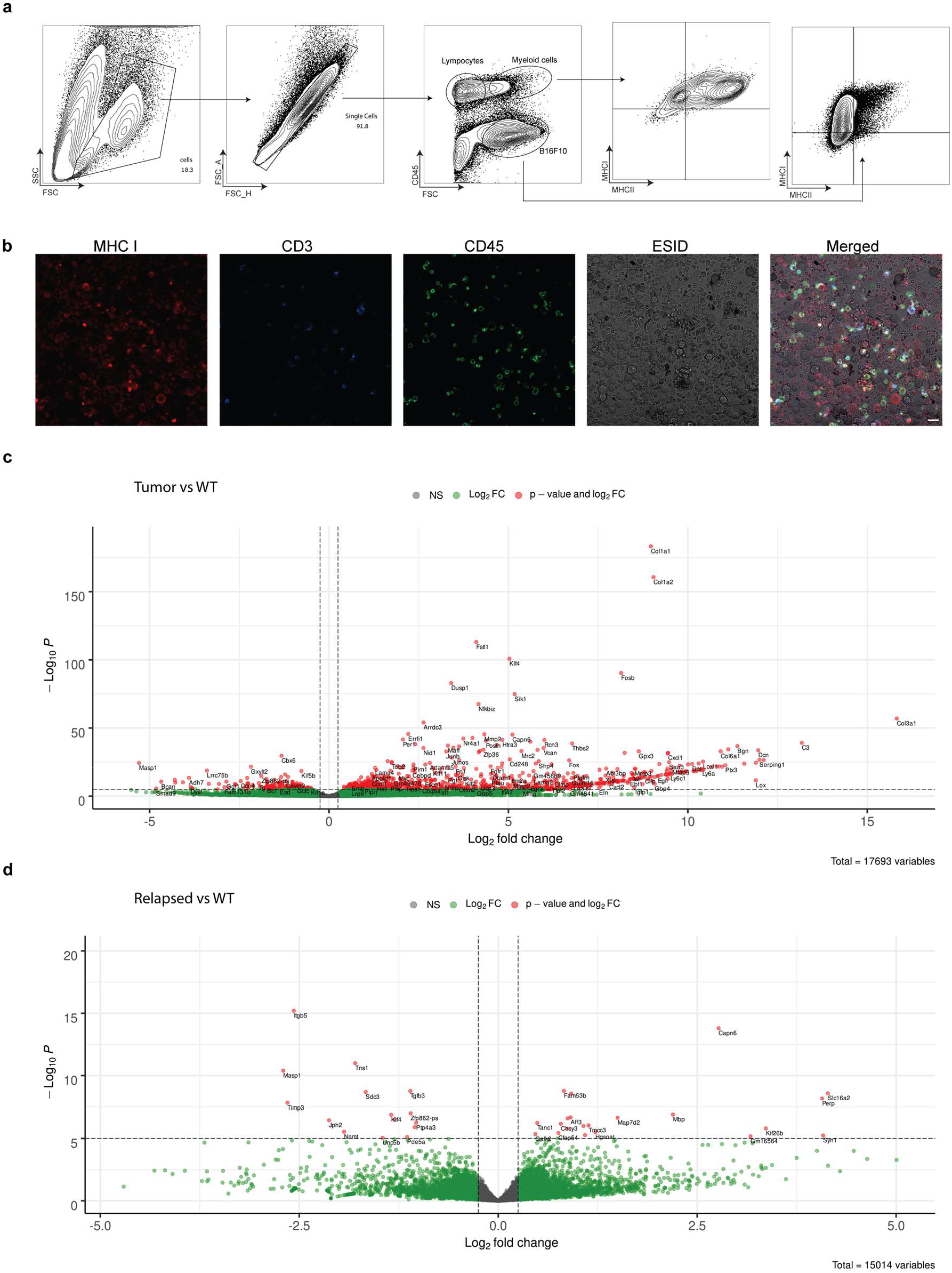
**a.** Gating strategy on immune and tumor cells following a second treatment with tumor-binding antibodies, TNFα, and anti-CD40 agonist. **b**. immunohistochemistry of total single cells obtained from tumors treated twice with tumorbinding antibodies, TNFα, and anti-CD40 agonist. **c. and d.** Volcano plots comparing gene expression in B16F10 tumor cells sorted from relapsed tumors *in vivo* and B16F10 cell line established from relapsed tumors and cultured *in vitro*. Scale bar = 20 μm.

**Extended Data Fig. 2:**
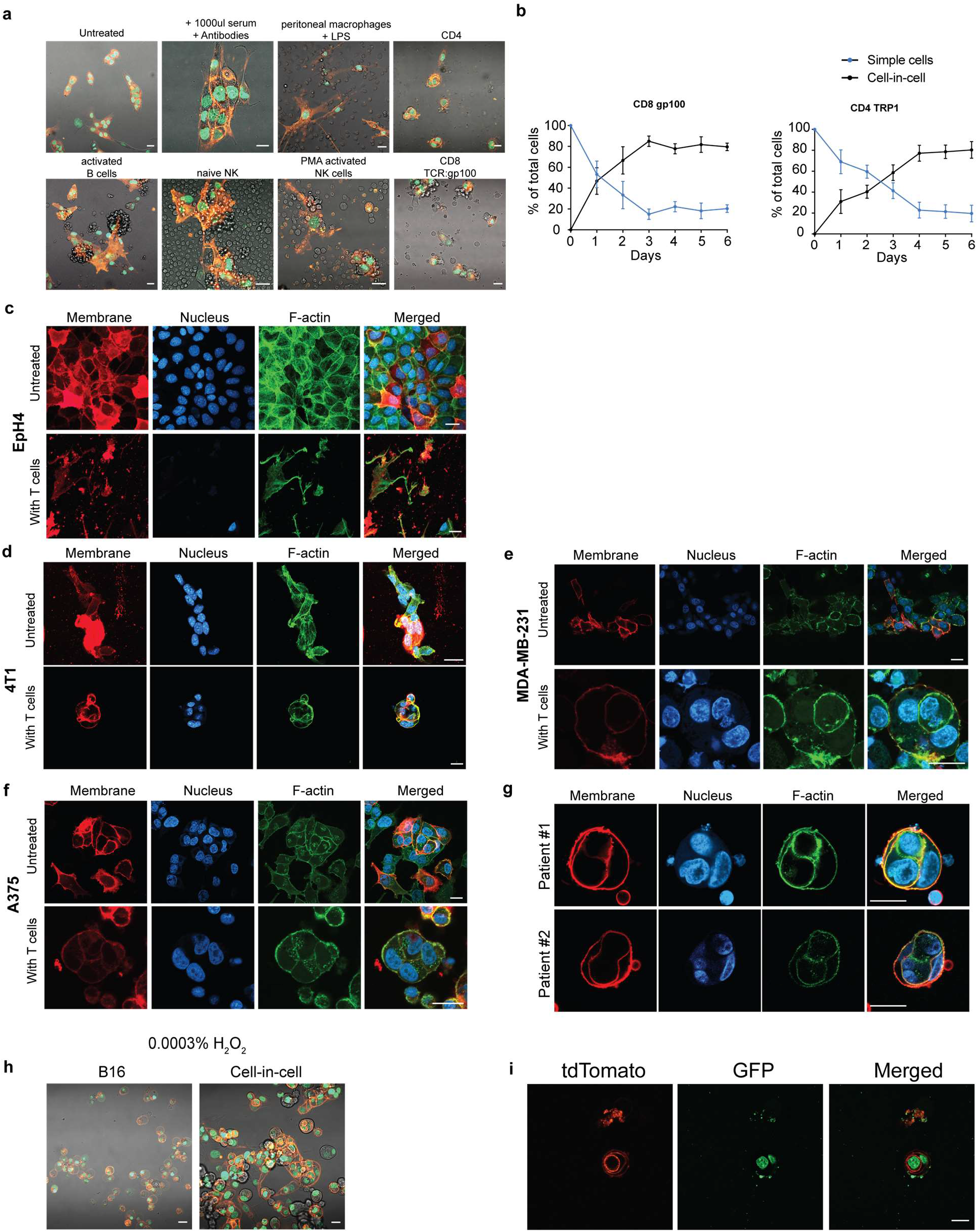
**a.** Representative images of B16F10 cells following incubation with different cells of the immune system. **b.** Mean percentage over time of cell-in-cell and simple cells from B16F10 tumor incubated with gp100-reactive CD8^+^ or TRP1-reactive CD4^+^ T cells. **c.** Representative images of breast epithelial non-transformed cells (EPH4) incubated with allogeneic T cells. **d.** Representative images of breast epithelial cancer cells (4T1) incubated with allogeneic T cells. **e.** Representative images of human breast cancer cell-line (MDA-MB-231) cells incubated with allogeneic T cells. **f.** Representative images of melanoma cell-line (A375) cells incubated with allogeneic T cells. **g.** Representative image of melanoma patients incubated with autologous T cells. **h.** Representative images of simple or cell-in-cell culture B16F10 incubated with H_2_O_2_. **i.** Representative images of B16F10 expressing H2B-GFP/MyrPalm-tdTomato incubated with CFSE stained T cell-derived lysosomes. Scale bars = 20 μm.

**Extended Data Fig. 3:**
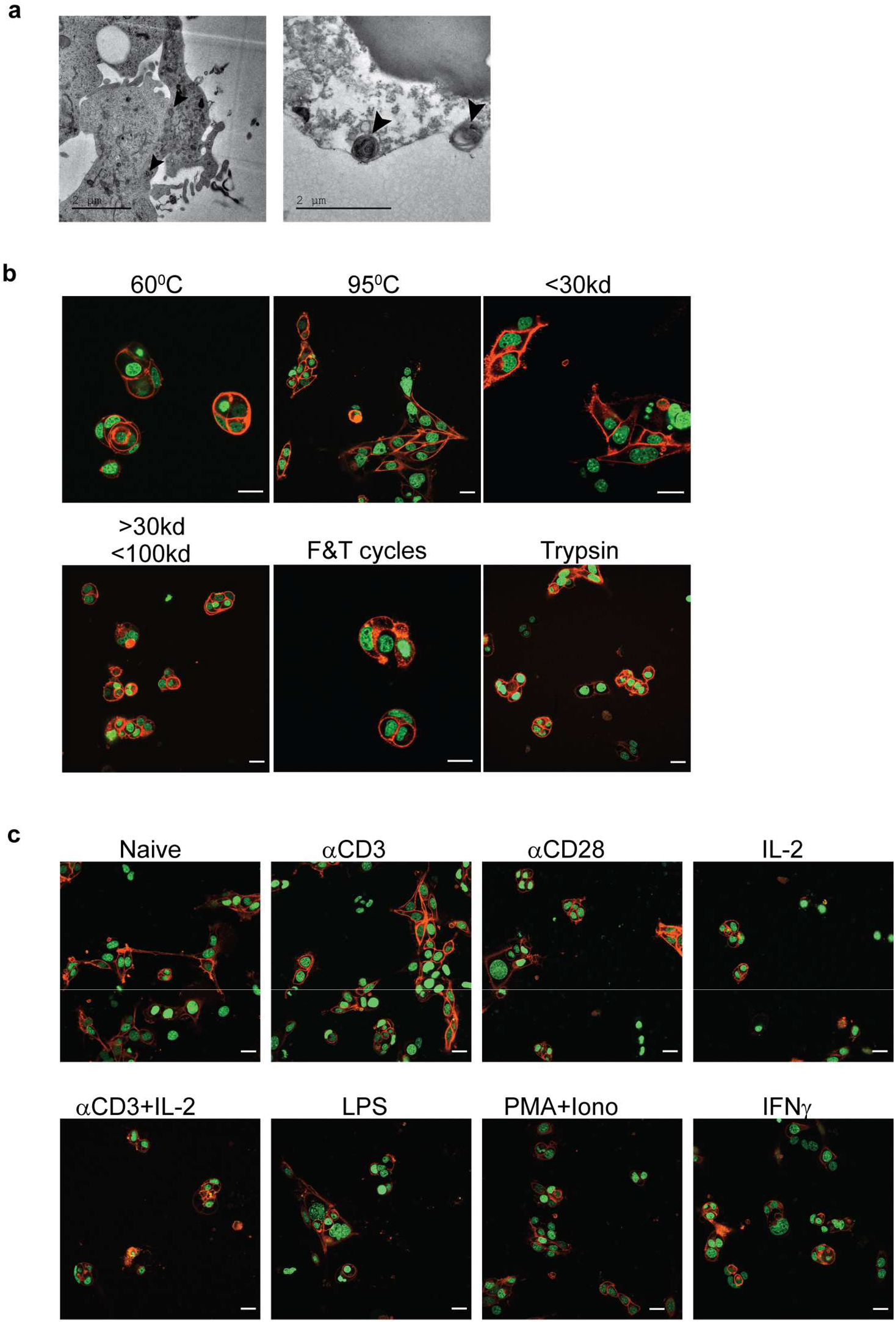
**a.** Single-layer transmitting electron microscopy of T cells incubated with B16F10. **b.** Representative images of B16F10 incubated overnight with T cell-derived lysosomes. **c.** Representative images of B16F10 incubated overnight with activated T cells. Scale bars = 20 μm (**b,c**).

**Extended Data Fig. 4:**
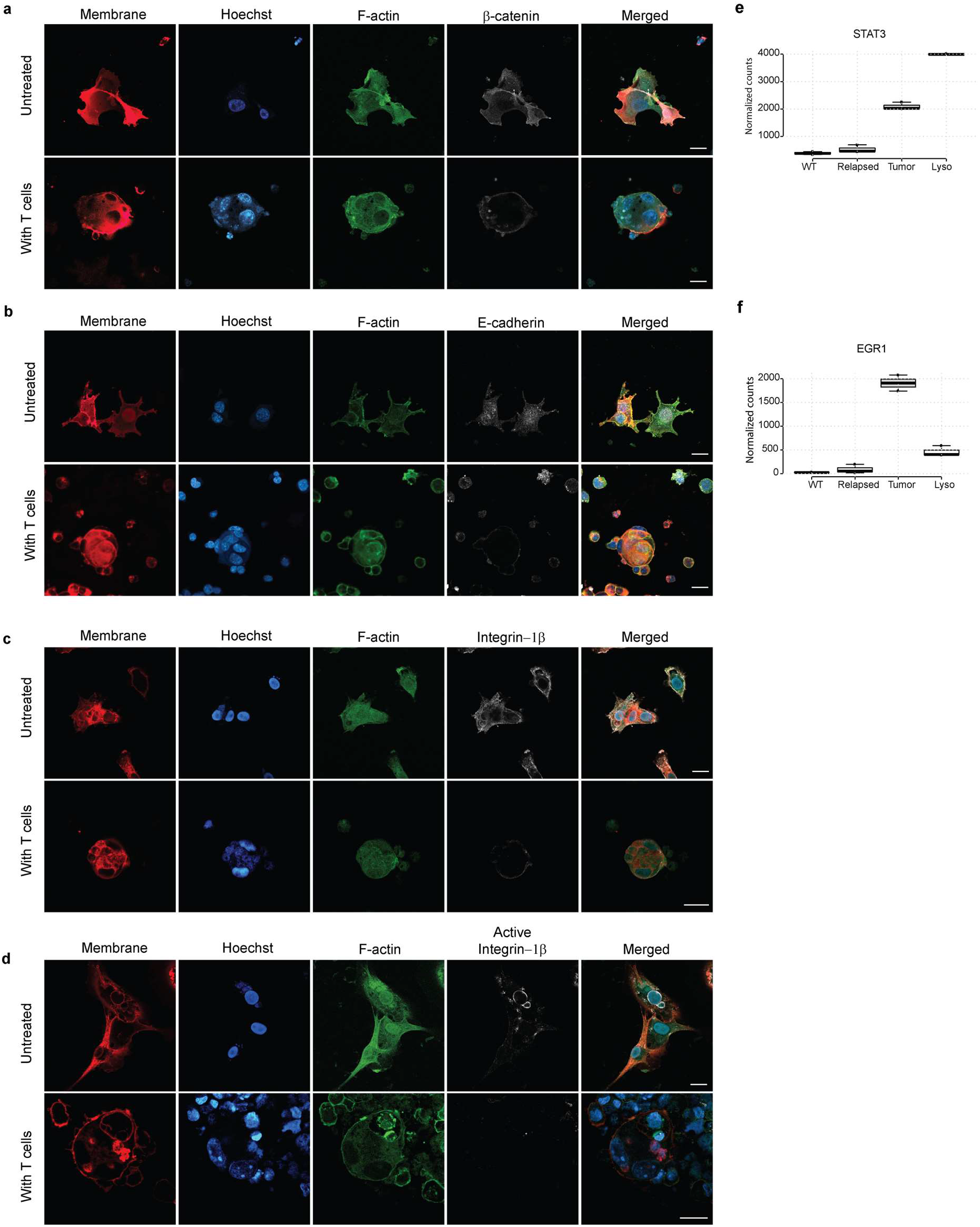
**a.** Representative images of β-catenin expression in B16F10 incubated overnight with gp100-reactive T cells. **b.** Representative images of E-cadherin expression in B16F10 incubated overnight with gp100-reactive T cells. **c.** Representative images of integrin-1 β expression on B16F10 incubated overnight with gp100-reactive T cells. **d.** Representative images of phosphorylated integrin-1β expression on B16F10 incubated overnight with gp100-reactive T cells. **e.** STAT3 expression level in B16F10 cells activated for 4 hours with T cell-derived lysosomes. **f**. EGR1 expression levels in B16F10 cells following 4 hours’ activation with T cell-derived lysosomes. Scale bars = 20 μm. Center lines show the medians; box limits indicate the 25th and 75th percentiles as determined by R software; whiskers extend 1.5 times the interquartile range from the 25th and 75th percentiles, outliers are represented by dots. n = 3 sample points.

## Extended Data Methods

### Mice

WT C57BL/6 and BALB/cOlaHsd mice were purchased from Envigo Israel. NSG (NOD/SCID-gamma KO mice) mice were a kind gift from Professor Michael Milyavsky at Tel Aviv University. All mice were housed in an American Association for the Accreditation of Laboratory Animal Care–accredited animal facility and maintained under specific pathogen-free conditions. Animal experiments were approved and conducted in accordance with Tel-Aviv University Laboratory Accreditation #01-16-095. Male and female 8-12 weeks old mice were used in all experiments.

### Cell lines

B16F10 cells (CRL-6475) and 4T1 (CRL-2539) cells were purchased directly from ATCC in January 2017 and used no later than passage four. A375 and MDA-MB-231 cell lines were a kind gift from Professor Tamar Geiger from Tel Aviv University. Both authentications were performed at the Genomics Core Facility of BioRap Technologies and the Rappaport Research Institute in Technion, Israel. Short tandem repeat profiles were determined using the Promega PowerPlex 16 HS Kit. HEK-293FT were purchased from Thermo Fisher Scientific in January 2017. EPH4 cells were a kind gift from professor Neta Erez from Tel Aviv University. Cells were cultured in DMEM (Gibco, Thermo Fisher Scientific) or RPMI1640 (Biological Industries) supplemented with 10% heat-inactivated FBS (Biological Industries), 2 mM l-glutamine, and 100 μg/mL penicillin/streptomycin (Gibco, Thermo Fisher Scientific) under standard conditions. Melan-A cells were purchased from Welcome Trust Functional Genomics Cell Bank in February 2019 and were supplemented with 200 nmol/L of phorbol 12-myristate 13-acetate (Santa Cruz Biotechnology). Cells were routinely tested for mycoplasma using a EZ-PCR Mycoplasma Test Kit (Biological Industries, Israel) according to the manufacturer’s instructions.

### Tumor models

For melanoma tumor studies, 2.5 × 10^5^ B16F10 cells suspended in 50 μL DMEM were injected s.c. into C57BL/6 mice above the right flank. For 4T1 triple-negative breast cancer model, 2 × 10^5^ 4T1 cells in 30 μL DMEM were injected into mammary fat pad number five fat pad number five. Tumor size was measured twice a week using calipers. Treatment was applied at different days post-injection. When tumors reached 120 mm^2^, the mice were sacrificed due to ethical considerations.

### Antibody-mediated immunotherapy

Tumor-bearing mice were injected intratumorally twice, two days apart, with 100 μg anti-CD40 (clone FGK4.5; BioXCell), 0.2 μg TNFα (BioLegend), and 200 μg/mouse anti-TRP1 antibody (clone TA99; BioXCell).

### Primary immune cell isolation

All tissue preparations were performed simultaneously from each individual mouse (after euthanasia by CO_2_ inhalation). Spleen was homogenized in 2% FBS and 5 μmol/L EDTA supplemented HBSS through 70 μm strainer (Thermo Fisher Scientific). Tumors were digested in RPMI 1640 with 2 mg/mL collagenase IV, 2,000 U/mL DNase I (both from Sigma Aldrich) for 40 minutes at 37°C with magnetic stirrers (200 rpm). Cells were then washed by centrifugation at 600 rcf for 5 min. at 4-8°C. Single cell suspension from tumor and spleens was applied of a Histopaque-1077 Hybri-Max™ (Sigma Aldrich) gradient and centrifuge for 15 minutes at 600 rcf.

For T-cell isolation, splenic mononuclear cells were incubated with anti-CD4 or anti-CD8 magnetic beads (MojoSort Nanobeads, BioLegend) according to the manufacturer’s instruction. For NK isolation, splenic mononuclear cells were incubated with anti-CD4, anti-CD8, anti-B220, anti-CD19 and anti-CD115 magnetic beads (MojoSort Nanobeads, BioLegend) according to the manufacturer’s instruction, and collected through negative selection.

Macrophages were isolated from peritoneal cavity of euthanized mice by washing and recollecting 5ml of HBSS twice, followed by centrifugation at 600 rcf.

Serum was acquired by pulling blood from venae cava of euthanized mice. Blood was incubated on ice for 30 minutes and centrifugation of at 400 rcf for 10 minutes. Serum was collected and recentrifuged at 21,000 rfc for 10 additional minutes.

### T-cell culture and expansion

T cells were cultured in RPMI 1640 supplemented with 1% Pen-Strep, 10% heat-inactivated FBS, 1% Sodium pyruvate, 1% MEM-Eagle non-essential amino acids, 1% Insulin-Transferrin-Selenium (PeproTech, Rocky Hill, NJ), 50 μM β-mercaptoethanol (Sigma-Aldrich, Merck, Israel) and, unless mentioned otherwise, were supplemented with 1,000 IU/mL recombinant murine IL-2 (PeproTech, Rocky Hill, NJ) on tissue culture plates pre-coated with 0.5 μg/mL anti-CD3 antibodies (clone 17A2, Biolegend).

### Tumor cell lentiviral transduction

For preparation of lentivirus, 1.3 × 10^6^ HEK-293FT cells were plated on a 6-well plate precoated with 200 μg/mL poly-L-lysine and let to adhere overnight. pLVX plasmids containing H2B-GFP, H2B-tdTomato, MyrPalm-tdTomato^50^, LifeAct-GFP, Wasabi or tdTomato under EF1 promoter together were mixed with psPAX2 (a gift from Didier Trono, Ecole Polytechnique Tédérale de Lausanne, Lausanne, Switzerland; Addgene plasmid 12260) and pCMV-VSV-G (a gift from Bob Weinberg, Massachusetts Institute of Technology, Cambridge, Massachusetts, USA; Addgene plasmid 8454) at a molar ratio of 3:2:1, and cells were transfected using Polyplus jetPRIME reagent (Polyplus Transfection). After 24 hours, medium was replaced with complete DMEM supplemented with 0.075% sodium bicarbonate. Medium-containing viruses were collected after 24 hours and 48 hours.

For tumor cell infection, virus-containing media were mixed with 100 μg/mL polybrene (Sigma-Aldrich) and added to B16F10 cells (1×10^6^ cells on 35 mm tissue culture) for 30 minutes at 37°C 5% CO_2_. Cells were centrifuged for 30 minutes at 37°C and 450 g. Then, 80% of the medium was replaced with complete DMEM supplemented with 0.075% sodium bicarbonate. After 3 days in culture, cells were sorted by FACSAriaIII. Cells were tested for mycoplasma, endotoxins, and bacterial contamination.

### T cells retrovirus Infection

HEK-293FT (Invitrogen) were plated on 10 cm culture plates and co-transfected with a 2:1 molar ratio of pMIGII and PCL-Eco plasmids (both were provided by Dario A.A. Vignali, University of Pittsburgh) using Polyplus jetPRIME reagent (Polyplus Transfection). After 24 hours, the medium was replaced with complete DMEM supplemented with 0.075% sodium bicarbonate. Media-containing viruses were collected after 24 hours and 48 hours and centrifuged for 1 hour at 100,000 g. The pellet was resuspended gently in 1 mL media and let to recover overnight at 4°C.

Prior to infection, splenic T cells were incubated on a plate precoated with anti-CD3 (0.5 μg/mL) in T cell medium containing high-dose IL-2 (1,000 IU/mL). Next, 0.3 mL concentrated retroviruses were added to every group of 2×10^6^ cells with 10 μg/mL polybrene. Cells were incubated for 30 minutes at 37°C in 5% CO_2_ and centrifuged at 37°C, 600 g, for one hour. Afterwards, 80% of medium was replaced and T cells were cultured for an additional three days in T cell media containing 1,000 IU IL-2. Transduction efficacy was assessed by FACS as the percentages of GFP-expressing cells.

### Adoptive T-cells Transfer

T cells were isolated from spleens of naïve mice as specified above and infected with pMIGII encoding TCR recognizing MHCI-gp100_25-33_^46^, or MHCI-TRP2_180-188_^47^, or MHCII-TRP1_113-126_^48^. Recipient tumo-bearing mice were sub-lethally irradiated at a single dose of 600 rad using cobalt source. After 24 hours, mice were injected i.v. twice, 3 days apart, with 1×10^6^ transduced T cells followed by i.p. injections of 300,000 IU of IL2 (PeproTech) for 4 consecutive days.

### Flow cytometry

For cell surface staining, monoclonal antibodies conjugated to FITC, PE, PE-Cy7, PE-Cy5.5, APC-Cy7, eFluor 650, or Pacific Blue and specific for the following antigens were used: CD11b (M1/70), Gr-1 (RB6-8C5), F4/80 (BM8), B220 (RA3-6B2), CD45 (A20), CD3 (17A2), CD4 (RM4-4), CD8 (53.6.7), MHCK^b^/K^d^ (28-8-6), CD115 (AFS98), I-Ab (M5/114.15.2), from BioLegend (San Diego, CA). Flow cytometry was performed on CytoFLEX (Beckman Coulter) and datasets were analyzed using FlowJo software (Tree Star, Inc.). All *in vivo* experiments to characterize tumor-infiltrating leukocytes were independently repeated at least 3 times with 3-5 mice per group. iTAg APC-labeled H-2K^b^-Trp-2_(SVYDFFVWL)_ and iTAg PE-labeled H-2D^b^-gp100_(EGSRNQDWL)_ tetramers were purchased from MBL international (Woburn, MA) and were used according to manufacturer’s instruction. Tetramers-staining experiments were repeated twice with 5 mice in each group.

### Confocal microscopy

For confocal microscopy imaging, cells were cultured on glass-bottom confocal plates (Cellvis). Cells were live-imaged or fixed and permeabilized with 2% PFA for 20 minutes. Fixed cultures were washed twice with PBS and stained for membranes with DiD membrane dye (Invitrogen), F-actin with Alexa Fluor™ 594 Phalloidin (Invitrogen) and nuclei with Hoechst 33342 (Fluka). For immunofluorescence, fixed cultures were blocked overnight with 5% BSA and stained with 1:100 or 1:200 diluted primary antibodies. Cells were then washed with PBS containing 1% BSA and stained with secondary antibody, diluted 1:100 or 1:200. We used the primary antibodies: anti-CD44 (clone IM7), anti-MHC class I (clone 28-8-6)(both from BioLegend), anti-β-catenin (clone D2U8Y), anti-phosphorylated STAT3 (clone D3A7), anti-active integrin 1β (polyclonal)(all from Cell Signaling Technology), anti-integrin 1β (clone 9EG7), anti-E-cadherin (clone 36)(both from BD Biosciences), anti-TRP2 (clone C-9, Santa Cruz Biotechnology), anti-gp100 (clone EP4863(2)) and anti-phosphorylated EGFR (clone EP38Y)(both from Abcam).

For frozen sections, tumor tissues were fixed in 4% paraformaldehyde for two hours and equilibrated in a 20% sucrose solution overnight. Tissues were then embedded in frozen tissue matrix (Scigen O.C.T. Compound Cryostat Embedding Medium, Thermo Fisher Scientific) and frizzed at −80°C. All specimens were imaged by ZEISS LSM800 (ZEISS) and analyzed by ZEN 2.3 (ZEISS) and Imagej.

### Electron microscopy

Scanning Electron Microscopy preparation was done as described by Fischer et al, with minor modifications^51^. Briefly, cells were fixed with warm 2.5% glutaraldehyde (GA) (EMS) in PBS for 60 minutes. Cells were then washed twice for 5 min in PBS, and twice with cacodylate buffer (0.1MCaCO, 5mMCaCl pH7.3) (Merck), post-fixed with 1% OsO_4_ for 60 minutes, washed 3 times in cacodylate buffer and then twice with H_2_O; Dehydration was done with increasing concentrations of reagents grade ethanol (2×5 minutes for 25%, 50%, 70%, 95% and 2×10 minutes for 100%), and was followed by coating with gold nanoparticles. Images were captured using GeminiSEM 300 (Carl Zeiss Microscopy, Jena, Thuringia, Germany).

For Transmitting Electron Microscopy cells were fixed in 2.5% Glutaraldehyde in PBS over night at 4°C. After several washings in PBS cells were post fixed in 1% OsO_4_ in PBS for 2h at 4°C. Dehydration was carried out in graded ethanol followed by embedding in Glycid ether. Thin sections were mounted on Formvar/Carbon coated grids, stained with uranyl acetate and lead citrate and examined in Jeol 1400 – Plus transmission electron microscope (Jeol, Japan). Images were captured using SIS Megaview III and iTEM the Tem imaging platform (Olympus).

### Incucyte assays

For long-term live cell imaging, B16F10 expressing H2B-TdTomato, H2B-GFP, and a mixture of both were sorted from immunotherapy-treated mice and plated in flat-bottom 96-well tissue culture (Corning) and placed in IncuCyte^®^ S3 Live Cell Analysis System (Essen BioScience). For killing assay, 1×10^4^ B16F10 cells expressing H2B-TdTomato were plated in flat-bottom 96-well tissue culture (10×10^3^ per well, Corning) and were let to adhere for several hours. Splenic CD8^+^ T cells infected with TCR against gp100, or TRP2 were added to culture at several ratios ranging from 1:1 to 10:1 E:T and placed in IncuCyte^®^ S3 Live Cell Analysis System (Essen BioScience). In other experiments, 1×10^4^ B16 cells expressing H2B-TdTomato were plated in flat-bottom 96-well tissue culture (Corning) and incubated in full DMEM medium containing either 1.5 mM doxorubicin, 3×10^-5^% H_2_O_2_ or 50 mg/ml cycloheximide for 92 hours. Confluence percentage and images were acquired and analyzed using IncuCyte^®^ Base Analysis Software (Essen BioScience).

### Neoantigen predication

Genomic DNA was extracted from 5×10^6^ B16F10 tumor cell lines using NucleoSpin^®^ Tissue (MACHEREY-NAGEL), according to manufacturer instructions. Whole exome sequencing was performed by NovoGene. Sequencing libraries were constructed using the Agilent SureSelect kit and sequence on an Illumina Novaseq 6000 sequencer. Sequencing reads were filtered using the AGeNT SureCall Trimmer (Agilent, v4.0.1) and quality was evaluated using FastQC v0.11.3. Alignment to the mouse genome (mm10, GRCm38.87) was performed using the Burrows-Wheeler Aligner (bwa v 0.7.12) using the bwa mem command with the following parameters: -M -t 4 -T 30. SAMtools was used to remove alignments with a MAPQ score less than 5, and Picard was used to remove duplicate reads. Sorting and indexing were performed using SAMtools. Variants were called using MuTect2 (GATK v 4.0.0.0) and filtered using FilterMutectCalls. Variants were annotated with SnpEff. Neoantigens were identified using MuPeXI v1.2.0^49^ with the following parameters: -s mouse -a H-2-Db, H-2-Kb -l 9-11. MuPeXI was used in conjunction with NetMHCpan v.4.0 and Variant Effect Predictor (VEP) v.87.27. Heatmaps were generated in R using the “pheatmap” package (v1.0.12).

### mRNA extraction and RNAseq analyses

To extract mRNA from tumor cells, 5×10^6^ B16F10 sorted from *in vivo* relapsed tumors, or from cell B16F10 cell lines established from relapsed tumors were collected. For preparation cell-incell culture induced by T cells-derived lysosomes, 1×10^6^ B16F10 cells were seeded on a low-adherence 10 cm plastic plate pre-coated with 200 μg/mL poly-L-lysine and were let to adhere for three hours. Concentrated T cell-derived lysosomes in DMEM were added to the culture, in 1:10 ratio for 4 hours. Cells were collected by washing with fresh media followed by 5 minutes centrifugation at 250 g. RNA was extracted from cells using NucleoSpin^®^ RNA (MACHEREY-NAGEL), according to manufacturer instructions. Sequencing Libraries were prepared using INCPM mRNA Seq. NextSeq 75 cycles reads were sequenced on 8 lanes of an Illumina nextseq. The output was ~18 million reads per sample.

For B16 RNAseq, sequencing adaptors were trimmed using fastp 0.19.6^52^ and aligned to the GRCm38 mouse assembly using STAR 2.7.1a^53^. Gene count matrix was analyzed for differential expression using DESeq2 1.24.0^54^ Enrichment analysis was performed on differentially expressed genes (adjusted p-value < 0.05) using clusterProfiler 3.12.0.^55^

For T cells RNAseq, reads were aligned using Kallisto^56^ to mouse genome version mm10, followed by further processing using the Bioconductor package DESeq2 1.24.0^54^ The data was normalized using TMM normalization, and differentially expressed genes were defined using the differential expression pipeline on the raw counts with a single call to the function DESeq (FDR-adjusted P value <0.05). Heatmap figures were generated using pheatmap package^57^ and clustered using Euclidian distance. Boxplots were generated using BoxPlotR.

### Cell-derived lysosomes and melanosomes isolation

For isolation of T-cell–derived lysosomes, 2×10^7^ CD8^+^ T cells were isolated from spleen and plated in 10 cm plate precoated with anti-CD3 antibodies and high-dose IL-2 (1000 IU/mL) for 48 hours in T cell media containing 10% FBS pre-centrifuged at 140,000 rpm for one h to deplete bovine-derived exosomes. Supernatants were collected and concentrated using Amicon^®^ Ultra (Merk) with 100kD filter. For isolation of NK cells-derived lysosomes, splenic 3×10^6^ NK cells isolated from naïve mice were plated in 12-well plate incubated with high-dose IL-2 for 48 hours. Supernatants were collected and concentrated using Amicon^®^ Ultra (Merk) with 100kD filter. In some experiments, lysosomes were pre-incubated 30 minutes in trypsin, fixed for 15 minutes in 2% PFA, heated for 50 minutes to 60°C or 5 minutes in 95°C, or were subjected to three freeze and thaw cycles. Lysosomes were added to B16F10 cultures for 24-48 hours incubation and the numbers of cell-in-cell structures out of total 200 cells were counted under fluorescent microscope.

For isolation of secreted melanosomes from B16F10, Melan A, 8×10^6^ cell were cultured in 10cm tissue culture plates and let to adhere overnight. Cells were incubated 48 h in DMEM supplemented with 10% exosome-free FBS. Supernatants was collected and centrifuged for 1 hour on glucose gradient, as previously described^58^.

### Cell-in-cell inhibition assays

For inhibition assays, B16F10 were plated in 4 Chamber glass-bottom confocal plates (Cellvis) 3,000 cells/chamber and let adhere for several hours. Cells were incubated for 24-48 hours in complete DMEM, with or without 50 μg/mL cycloheximide, 120 mM 5,6-dichloro-1-beta-D-ribofuranosylbenzimidazole (DRB) (Sigma-Aldrich), 20 nM Latrunculin B (Lat-B, Sigma-Aldrich), 2 μM Stattic (Sigma-Aldrich), 4 μM Gefitinib, 5 μM XAV-939 (Sigma-Aldrich) or the blocking antibodies anti-E-cadherin (clone 36) and anti-N-cadherin (clone 13A9). In addition, gp100-reactive T cells (E:T ratio of 5:1), or T cells-derived lysosomes (10-50 μL of concentrated lysosomes per well) were added to the culture. Cell-in-cell structures were counted under fluorescent microscope for cell-in-cell per 200 cells, three times for each chamber.

### Cell-in-cell counts

For cell-in-cell count, 3,000 tumor cells per one cm^2^ were plated in a 4 Chamber glass-bottom confocal plates (Cellvis) and let adhere for several hours. Cells were then incubated overnight to 48 hours with immune cells, or exosomes and counted under fluorescent microscope. Percentages of cell-in-cells structures were calculated by counting 200 cells in three different chambers, three fields in each chamber.

### Study approval

All animal protocols were approved by the Stanford University Institutional Animal Care and Use Committee under protocol: 01-16-095. The Tel Aviv University Institutional Review Board approved the human subject protocols, and informed consent was obtained from all subjects prior to participation in the study.

## Acknowledgments

The authors would like to deeply thanks Rabin Medical Center Institutional Tissue Bank and especially Dr. Adva Levi-Barda, for their invaluable support of this research. We would also like to thank Dr. Vered Holdengreber from the Faculty of Life Sciences, Tel Aviv University and Dr. Gal Radovsky from Tel Aviv University Center for Nano Science and Nano Technology for their help with electron microscopy imaging.

## Author contributions

AG has conducted the majority of the experiments and helped to design them. NSM, DR, and LFY have helped to conduct the experiments. AM has analyzed T cells RNAseq data. CL, JC, and KP have provided lentiviral constracts and viral particals for labeling. NS and GS have analyzed B16 cells RNAseq data. MF and OZ have provided biological materials from human melanoma patients. RD has provided TCR:TRP2 construct. NRF has analyzed WES data and neoantigan analyzis. PR and YC have designed and conducted the experiments and have written the manuscript.

## Statistics

For time course experiments, significance was calculated using the nonparametric two-way ANOVA with Tukey’s correction for multiple hypotheses. In some cases, Bonferroni-Sidak post-test was performed after two-way ANOVA. For two groups analysis, one-way ANOVA with Dunn’s test was performed. The results were analyzed by Prism (GraphPad Software, Inc.). All statistical analyses were performed in Prism (GraphPad Software, Inc.). All experiments were performed at least 3 times with at least three replications.

## References

1 Pages, F. et al. Effector memory T cells, early metastasis, and survival in colorectal cancer. N Engl J Med 353, 2654–2666, doi:10.1056/NEJMoa051424 (2005).

2 Galon, J. et al. Type, density, and location of immune cells within human colorectal tumors predict clinical outcome. Science 313, 1960–1964, doi:10.1126/science.1129139 (2006).

3 Fridman, W. H., Pages, F., Sautes-Fridman, C. & Galon, J. The immune contexture in human tumours: impact on clinical outcome. Nat Rev Cancer 12, 298–306, doi:10.1038/nrc3245 (2012).

4 Restifo, N. P., Dudley, M. E. & Rosenberg, S. A. Adoptive immunotherapy for cancer: harnessing the T cell response. Nat Rev Immunol 12, 269–281, doi:10.1038/nri3191 (2012).

5 Rosenberg, S. A. Decade in review-cancer immunotherapy: entering the mainstream of cancer treatment. Nat Rev Clin Oncol 11, 630–632, doi:10.1038/nrclinonc.2014.174 nrclinonc.2014.174 [pii] (2014).

6 Gross, G., Gorochov, G., Waks, T. & Eshhar, Z. Generation of effector T cells expressing chimeric T cell receptor with antibody type-specificity. Transplant Proc 21, 127–130 (1989).

7 Kalos, M. et al. T cells with chimeric antigen receptors have potent antitumor effects and can establish memory in patients with advanced leukemia. Sci Transl Med 3, 95ra73, doi:10.1126/scitranslmed.3002842 3/95/95ra73 [pii] (2011).

8 Pardoll, D. M. The blockade of immune checkpoints in cancer immunotherapy. Nat Rev Cancer 12, 252–264, doi:10.1038/nrc3239 nrc3239 [pii] (2012).

9 Hodi, F. S. et al. Improved survival with ipilimumab in patients with metastatic melanoma. The New England journal of medicine 363, 711–723, doi:10.1056/NEJMoa1003466 (2010).

10 Salati, M., Baldessari, C., Cerbelli, B. & Botticelli, A. Nivolumab in pretreated nonsmall cell lung cancer: continuing the immunolution. Transl Lung Cancer Res 7, S91–S94, doi:10.21037/tlcr.2018.01.14 (2018).

11 Gide, T. N., Wilmott, J. S., Scolyer, R. A. & Long, G. V. Primary and Acquired Resistance to Immune Checkpoint Inhibitors in Metastatic Melanoma. Clin Cancer Res 24, 1260–1270, doi:10.1158/1078-0432.CCR-17-2267 (2018).

12 Gauci, M. L. et al. Long-Term Survival in Patients Responding to Anti-PD-1/PD-L1 Therapy and Disease Outcome upon Treatment Discontinuation. Clin Cancer Res 25, 946–956, doi:10.1158/1078-0432.CCR-18-0793 (2019).

13 Vogelstein, B. et al. Cancer genome landscapes. Science 339, 1546–1558, doi:10.1126/science.1235122 339/6127/1546 [pii] (2013).

14 Schumacher, T. N. & Schreiber, R. D. Neoantigens in cancer immunotherapy. Science 348, 69–74, doi:10.1126/science.aaa4971 348/6230/69 [pii] (2015).

15 Khong, H. T. & Restifo, N. P. Natural selection of tumor variants in the generation of “tumor escape” phenotypes. Nature Immunology 3, 999–1005, doi:10.1038/ni1102-999 (2002).

16 Dunn, G. P., Bruce, A. T., Ikeda, H., Old, L. J. & Schreiber, R. D. Cancer immunoediting: from immunosurveillance to tumor escape. Nature Immunology 3, 991–998, doi:10.1038/ni1102-991 (2002).

17 Gajewski, T. F., Schreiber, H. & Fu, Y.-X. Innate and adaptive immune cells in the tumor microenvironment. Nature Immunology 14, 1014, doi:10.1038/ni.2703 (2013).

18 Zaretsky, J. M. et al. Mutations Associated with Acquired Resistance to PD-1 Blockade in Melanoma. N Engl J Med 375, 819–829, doi:10.1056/NEJMoa1604958 (2016).

19 Sade-Feldman, M. et al. Resistance to checkpoint blockade therapy through inactivation of antigen presentation. Nat Commun 8, 1136, doi:10.1038/s41467-017-01062-w (2017).

20 Peng, W. et al. Loss of PTEN Promotes Resistance to T Cell-Mediated Immunotherapy. Cancer Discov 6, 202–216, doi:10.1158/2159-8290.CD-15-0283 (2016).

21 Yates, L. R. et al. Genomic Evolution of Breast Cancer Metastasis and Relapse. Cancer Cell 32, 169–184 e167, doi:10.1016/j.ccell.2017.07.005 (2017).

22 Hao, J. J. et al. Spatial intratumoral heterogeneity and temporal clonal evolution in esophageal squamous cell carcinoma. Nat Genet 48, 1500–1507, doi:10.1038/ng.3683 (2016).

23 Gundem, G. et al. The evolutionary history of lethal metastatic prostate cancer. Nature 520, 353–357, doi:10.1038/nature14347 (2015).

24 Carmi, Y. et al. Allogeneic IgG combined with dendritic cell stimuli induce antitumour T-cell immunity. Nature 521, 99–104, doi:10.1038/nature14424 nature14424 [pii] (2015).

25 Overholtzer, M. et al. A nonapoptotic cell death process, entosis, that occurs by cellin-cell invasion. Cell 131, 966–979, doi:10.1016/j.cell.2007.10.040 (2007).

26 Mittal, D., Gubin, M. M., Schreiber, R. D. & Smyth, M. J. New insights into cancer immunoediting and its three component phases--elimination, equilibrium and escape. Current opinion in immunology 27, 16–25, doi:10.1016/j.coi.2014.01.004 (2014).

27 Matsushita, H. et al. Cancer exome analysis reveals a T-cell-dependent mechanism of cancer immunoediting. Nature 482, 400–404, doi:10.1038/nature10755 (2012).

28 DuPage, M., Mazumdar, C., Schmidt, L. M., Cheung, A. F. & Jacks, T. Expression of tumour-specific antigens underlies cancer immunoediting. Nature 482, 405–409, doi:10.1038/nature10803 (2012).

29 Verdegaal, E. M. et al. Neoantigen landscape dynamics during human melanoma-T cell interactions. Nature 536, 91–95, doi:10.1038/nature18945 (2016).

30 von Boehmer, L. et al. NY-ESO-1-specific immunological pressure and escape in a patient with metastatic melanoma. Cancer Immun 13, 12 (2013).

31 Tran, E. et al. T-Cell Transfer Therapy Targeting Mutant KRAS in Cancer. N Engl J Med 375, 2255–2262, doi:10.1056/NEJMoa1609279 (2016).

32 Tran, E., Robbins, P. F. & Rosenberg, S. A. ‘Final common pathway’ of human cancer immunotherapy: targeting random somatic mutations. Nat Immunol 18, 255–262, doi:10.1038/ni.3682 (2017).

33 Karanikas, V. et al. High frequency of cytolytic T lymphocytes directed against a tumor-specific mutated antigen detectable with HLA tetramers in the blood of a lung carcinoma patient with long survival. Cancer Res 61, 3718–3724 (2001).

34 Prickett, T. D. et al. Durable Complete Response from Metastatic Melanoma after Transfer of Autologous T Cells Recognizing 10 Mutated Tumor Antigens. Cancer Immunol Res 4, 669–678, doi:10.1158/2326-6066.CIR-15-0215 (2016).

35 Birnbaum, M. E. et al. Deconstructing the peptide-MHC specificity of T cell recognition. Cell 157, 1073–1087, doi:10.1016/j.cell.2014.03.047 (2014).

36 Wooldridge, L. et al. A single autoimmune T cell receptor recognizes more than a million different peptides. J Biol Chem 287, 1168–1177, doi:10.1074/jbc.M111.289488 (2012).

37 Bentzen, A. K. et al. T cell receptor fingerprinting enables in-depth characterization of the interactions governing recognition of peptide-MHC complexes. Nat Biotechnol, doi:10.1038/nbt.4303 (2018).

38 Krishna, S. & Overholtzer, M. Mechanisms and consequences of entosis. Cell Mol Life Sci 73, 2379–2386, doi:10.1007/s00018-016-2207-0 (2016).

39 Tonnessen-Murray, C. A. et al. Chemotherapy-induced senescent cancer cells engulf other cells to enhance their survival. J Cell Biol 218, 3827–3844, doi:10.1083/jcb.201904051 (2019).

40 Hamann, J. C. et al. Entosis Is Induced by Glucose Starvation. Cell Rep 20, 201–210, doi:10.1016/j.celrep.2017.06.037 (2017).

41 Durgan, J. et al. Mitosis can drive cell cannibalism through entosis. Elife 6, doi:10.7554/eLife.27134 (2017).

42 Overwijk, W. W. et al. gp100/pmel 17 is a murine tumor rejection antigen: induction of “self’-reactive, tumoricidal T cells using high-affinity, altered peptide ligand. J Exp Med 188, 277–286, doi:10.1084/jem.188.2.277 (1998).

43 Bendle, G. M. et al. Lethal graft-versus-host disease in mouse models of T cell receptor gene therapy. Nat Med 16, 565–570, 561p following 570, doi:10.1038/nm.2128 (2010).

44 Muranski, P. et al. Tumor-specific Th17-polarized cells eradicate large established melanoma. Blood 112, 362–373, doi:10.1182/blood-2007-11-120998 (2008).

45 Bjerregaard, A. M., Nielsen, M., Hadrup, S. R., Szallasi, Z. & Eklund, A. C. MuPeXI: prediction of neo-epitopes from tumor sequencing data. Cancer Immunol Immunother 66, 1123–1130, doi:10.1007/s00262-017-2001-3 (2017).

## Extended Reference

7 Kalos, M. et al. T cells with chimeric antigen receptors have potent antitumor effects and can establish memory in patients with advanced leukemia. Sci TranslMed 3, 95ra73, doi:10.1126/scitranslmed.3002842 3/95/95ra73 [pii] (2011).

33 Straetemans, T. et al. Recurrence of melanoma following T cell treatment: continued antigen expression in a tumor that evades T cell recruitment. Mol Ther 23, 396–406, doi:10.1038/mt.2014.215 (2015).

34 Karanikas, V. et al. High frequency of cytolytic T lymphocytes directed against a tumor-specific mutated antigen detectable with HLA tetramers in the blood of a lung carcinoma patient with long survival. Cancer Res 61, 3718–3724 (2001).

35 Prickett, T. D. et al. Durable Complete Response from Metastatic Melanoma after Transfer of Autologous T Cells Recognizing 10 Mutated Tumor Antigens. Cancer Immunol Res 4, 669–678, doi:10.1158/2326-6066.CIR-15-0215 (2016).

36 Birnbaum, M. E. et al. Deconstructing the peptide-MHC specificity of T cell recognition. Cell 157, 1073–1087, doi:10.1016/j.cell.2014.03.047 (2014).

37 Wooldridge, L. et al. A single autoimmune T cell receptor recognizes more than a million different peptides. J Biol Chem 287, 1168–1177, doi:10.1074/jbc.M111.289488 (2012).

38 Bentzen, A. K. et al. T cell receptor fingerprinting enables in-depth characterization of the interactions governing recognition of peptide-MHC complexes. Nat Biotechnol, doi:10.1038/nbt.4303 (2018).

39 Thorsson, V. et al. The Immune Landscape of Cancer. Immunity 48, 812–830 e814, doi:10.1016/j.immuni.2018.03.023 (2018).

40 Joyce, J. A. & Fearon, D. T. T cell exclusion, immune privilege, and the tumor microenvironment. Science 348, 74–80, doi:10.1126/science.aaa6204 348/6230/74 [pii] (2015).

41 Gabrilovich, D. I., Ostrand-Rosenberg, S. & Bronte, V. Coordinated regulation of myeloid cells by tumours. Nature reviews. Immunology 12, 253–268, doi:10.1038/nri3175 (2012).

42 Krishna, S. & Overholtzer, M. Mechanisms and consequences of entosis. Cell Mol Life Sci 73, 2379–2386, doi:10.1007/s00018-016-2207-0 (2016).

43 Tonnessen-Murray, C. A. et al. Chemotherapy-induced senescent cancer cells engulf other cells to enhance their survival. J Cell Biol 218, 3827–3844, doi:10.1083/jcb.201904051 (2019).

44 Hamann, J. C. et al. Entosis Is Induced by Glucose Starvation. Cell Rep 20, 201–210, doi:10.1016/j.celrep.2017.06.037 (2017).

45 Durgan, J. et al. Mitosis can drive cell cannibalism through entosis. Elife 6, doi:10.7554/eLife.27134 (2017).

46 Overwijk, W. W. et al. gp100/pmel 17 is a murine tumor rejection antigen: induction of “self’-reactive, tumoricidal T cells using high-affinity, altered peptide ligand. J Exp Med 188, 277–286, doi:10.1084/jem.188.2.277 (1998).

47 Bendle, G. M. et al. Lethal graft-versus-host disease in mouse models of T cell receptor gene therapy. Nat Med 16, 565–570, 561p following 570, doi:10.1038/nm.2128 (2010).

48 Muranski, P. et al. Tumor-specific Th17-polarized cells eradicate large established melanoma. Blood 112, 362–373, doi:10.1182/blood-2007-11-120998 (2008).

49 Bjerregaard, A. M., Nielsen, M., Hadrup, S. R., Szallasi, Z. & Eklund, A. C. MuPeXI: prediction of neo-epitopes from tumor sequencing data. Cancer Immunol Immunother 66, 1123–1130, doi:10.1007/s00262-017-2001-3 (2017).

50 Zacharias, D. A., Violin, J. D., Newton, A. C. & Tsien, R. Y. Partitioning of lipid-modified monomeric GFPs into membrane microdomains of live cells. Science 296, 913–916, doi:10.1126/science.1068539 (2002).

51 Fischer, E. R., Hansen, B. T., Nair, V., Hoyt, F. H. & Dorward, D. W. Scanning electron microscopy. Curr Protoc Microbiol Chapter 2, Unit 2B 2, doi:10.1002/9780471729259.mc02b02s25 (2012).

52 Chen, S., Zhou, Y., Chen, Y. & Gu, J. fastp: an ultra-fast all-in-one FASTQ preprocessor. Bioinformatics 34, i884–i890, doi:10.1093/bioinformatics/bty560 (2018).

53 Dobin, A. et al. STAR: ultrafast universal RNA-seq aligner. Bioinformatics 29, 15–21, doi:10.1093/bioinformatics/bts635 (2012).

54 Love, M. I., Huber, W. & Anders, S. Moderated estimation of fold change and dispersion for RNA-seq data with DESeq2. Genome Biology 15, 550, doi:10.1186/s13059-014-0550-8 (2014).

55 Yu, G., Wang, L.-G., Han, Y. & He, Q.-Y. clusterProfiler: an R package for comparing biological themes among gene clusters. Omics: a journal of integrative biology 16, 284–287, doi:10.1089/omi.2011.0118 (2012).

56 Bray, N. L., Pimentel, H., Melsted, P. & Pachter, L. Near-optimal probabilistic RNAseq quantification. Nature Biotechnology 34, 525–527, doi:10.1038/nbt.3519 (2016).

57 Kolde, R. & Vilo, J. GOsummaries: an R Package for Visual Functional Annotation of Experimental Data. F1000Research 4, 574–574, doi:10.12688/f1000research.6925.1 (2015).

58 Santana-Magal, N. et al. Melanoma-Secreted Lysosomes Trigger Monocyte-Derived Dendritic Cell Apoptosis and Limit Cancer Immunotherapy. Cancer Res 80, 1942–1956, doi:10.1158/0008-5472.CAN-19-2944 (2020).

